# Quantifying negative selection in human 3’ UTRs uncovers constrained targets of RNA-binding proteins

**DOI:** 10.1101/2022.11.30.518628

**Authors:** Scott D. Findlay, Lindsay Romo, Christopher B. Burge

## Abstract

Many non-coding variants associated with phenotypes occur in 3’ untranslated regions (3’ UTRs) and may affect interactions with RNA-binding proteins (RBPs) to regulate post-transcriptional gene expression. However, identifying functional 3’ UTR variants has proven difficult. We used allele frequencies from the Genome Aggregation Database (gnomAD) to identify classes of 3’ UTR variants under strong negative selection in humans. We developed intergenic mutability-adjusted proportion singleton (iMAPS), a generalized measure related to MAPS, to quantify negative selection in non-coding regions. This approach, in conjunction with *in vitro* and *in vivo* binding data, identifies precise RBP binding sites, miRNA target sites, and polyadenylation signals (PASs) under strong selection. For each class of sites, we identified thousands of gnomAD variants under selection comparable to missense coding variants, and found that sites in core 3’ UTR regions upstream of the most-used PAS are under strongest selection. Together, this work improves our understanding of selection on human genes and validates approaches for interpreting genetic variants in human 3’ UTRs.

## Introduction

Since the sequencing of the human genome, identifying functional genetic variants that influence human phenotypes including disease has been a central goal. For variants lying in protein-coding exons, the genetic code aids greatly in interpretation. However, the vast majority of candidate causal variants emerging from genome-wide association studies (GWAS) lie outside of protein-coding regions (Hindorff et al. 2009; Gusev et al. 2014; Maurano et al. 2012), and likely impact a variety of regulatory elements, making interpretation much more challenging (Dunham et al. 2012; Abascal et al. 2020).

Since transcription is a major point of regulation for gene expression, much of the search for functional non-coding variants has focused on transcriptional regulation and regions upstream of the coding sequence (Fulco et al. 2019; Wright et al. 2021; Deplancke et al. 2016). Other regions such as 3’ untranslated regions (3’ UTRs) have been less explored, despite playing major roles in post-transcriptional regulation via processes such as cleavage and polyadenylation, mRNA stability, and translation (Mayya and Duchaine 2019). 3’ UTRs explained GWAS genotyped SNP heritability at a rate 5-fold higher than expected (Finucane et al. 2015), and expression quantitative trait loci (eQTLs) were more enriched (> 2-fold) in 3’ UTRs than any other non-coding annotation analyzed (Consortium 2020), demonstrating an abundance of impactful genetic variation in 3’ UTRs.

Mechanistically, interactions between RNA-binding proteins (RBPs) and their target RNAs lie at the heart of virtually all post-transcriptional gene regulation in 3’ UTRs. As examples, Argonaute proteins guided by cellular microRNAs (miRNAs) bind most mRNAs to repress expression (Bartel 2009), cleavage and polyadenylation specificity factors (CPSFs) bind polyadenylation signals (PASs) to define 3’ ends of transcripts (Sun et al. 2018; Chan et al. 2014; Schönemann et al. 2014), and Pumilio family proteins (PUM1/PUM2) and AU-rich element (ARE) binding proteins such as TIA1 destabilize bound transcripts (Meyer et al. 2018; HafezQorani et al. 2016; Etten et al. 2012; Wolfe et al. 2020). Thus, the collection of RBP binding sites in 3’ UTRs constitutes a set of non-coding elements enriched for regulatory activity.

Recent large-scale efforts using techniques such as enhanced crosslinking and immunoprecipitation (eCLIP) have characterized RBP binding sites throughout the transcriptome (Van Nostrand et al. 2020), providing a basis for the discovery of allele-specific RNA-RBP binding events (Yang et al. 2019; Feng et al. 2019). However, these studies are inherently limited to the small number of variants that are heterozygous in the cell lines used, and little work has been done to more broadly assess the evolutionary pressures faced by all variants that modulate RBP-RNA interactions. The premise of such an approach is that the number of times variant alleles are observed in human populations will be lower within constrained functional regulatory elements in the genome, leaving a signature of negative/purifying selection during the “natural experiment” of human evolution. Such efforts were initially limited in resolution by the number of genomes with genome-wide and deep sequencing data available (Auton et al. 2015; Ward and Kellis 2012). More recently, projects such as the genome aggregation database (gnomAD) and UK Biobank have cataloged genetic variation from tens of thousands of whole genomes (Karczewski et al. 2020; Halldorsson et al. 2022). The gnomAD Consortium also developed the Mutability-Adjusted Proportion Singleton (MAPS) metric that summarizes the allele frequency spectrum across collections of variants to quantify negative selection. This metric improved on previous measures by capturing non-selective but well known forces impacting allele frequency spectra, such as differential mutability, that can greatly confound evaluation of negative selection (Lek et al. 2016; Karczewski et al. 2020; Harpak et al. 2016; Carlson et al. 2018; Rands et al. 2014).

While this type of approach has been applied to identify genetic variation under strong selection in non-coding regions such as 5’ UTRs (Whiffin et al. 2020) and introns at splice sites (Blakes et al. 2022; Lord et al. 2019), it has not yet been applied to 3’ UTRs, suggesting that signals of negative selection may be challenging to uncover in these regions. Instead, the few instances where negative selection in 3’ UTRs has been inferred have been limited in scope, did not adjust for mutability, and were secondary to other efforts (Zhang et al. 2020; Park et al. 2021; Kainov et al. 2016).

In this work, we detail patterns of negative selection across diverse classes of regulatory elements in human 3’ UTRs. We introduce the intergenic MAPS (iMAPS) approach that is well-suited to detect signals of negative selection in non-coding regions of the transcriptome. Using this method, we confirm a major role for RBP-RNA interactions in shaping the 3’ UTR regulatory landscape by describing numerous classes of genetic variants that are under strong selection, influence transcript levels, and can improve interpretation of non-coding genetic variants

## Results

We used the allele frequency spectrum to quantify negative selection in 3’ UTRs, where a shift toward rare variants is indicative of negative selection. Previous work quantifying negative selection in broadly-defined genomic regions has typically relied on calibrating the allele frequencies of variants of interest to synonymous coding variants. Variants were matched based on base change and flanking dinucleotide contexts in an effort to control for the influence of mutability (and other non-selective forces) on the allele frequency spectrum (Karczewski et al. 2020), while other work has not explicitly calibrated at this resolution (Zhang et al. 2020). We reasoned that these approaches may not be ideal for assessing more specific classes of non-coding regulatory variation where sequence composition is often less complex (Dominguez et al. 2018) and the magnitude of negative selection is likely more modest. Furthermore, some synonymous variation is clearly under selection in connection to RNA splicing, mRNA stability, etc., so calibrating to a more neutrally evolving class of variation is desirable. We developed a rigorous method to better account for the influence of mutability on the allele frequency spectrum by 1) calibrating to intergenic regions, 2) considering additional sequence context, and 3) considering the impact of transcription on DNA repair and mutation (Fig. 1A) for the reasons described below.

**Fig. 1:**
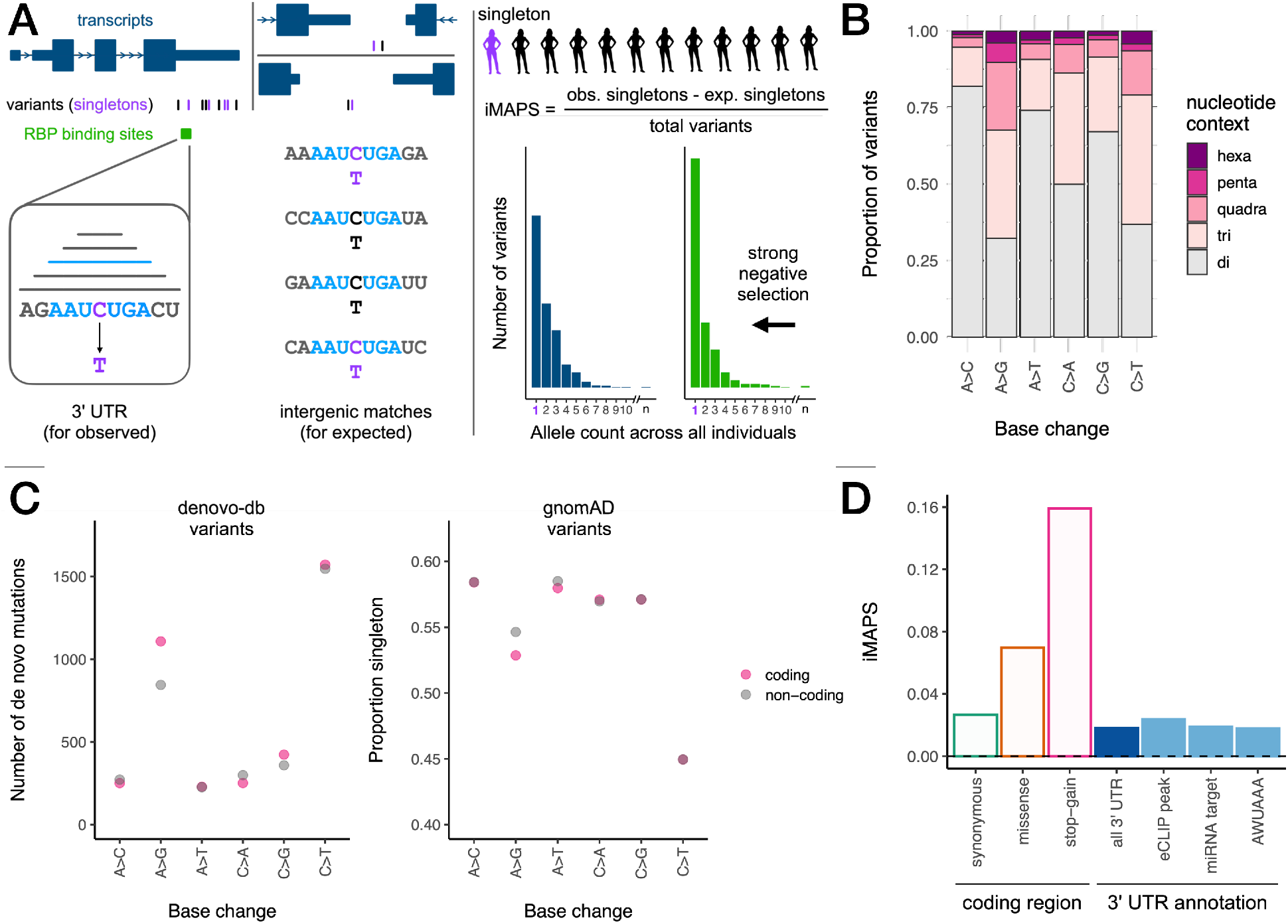
Using iMAPS to quantify negative selection in 3’ UTRs. A) Schematic of iMAPS approach. Left: each 3’ UTR variant is matched to multiple intergenic variants with the same base change and flanking sequence context (ranging from dinucleotide to hexanucleotide). Right: Top: the number of singletons (variant alleles detected once across all individuals in gnomAD) is used to calculate iMAPS. “obs” = observed, exp. = expected. Bottom: sets of variants under strong selection (green) have an excess of singletons and thus higher iMAPS scores. B) The proportion of 3’ UTR variants for which considering extended nucleotide contexts when matching to intergenic variants affects the expected singleton rate. C) Left: A>G/T>C mutations have the highest strand bias in de novo mutation rate relative to all other base changes. Right: accordingly, the proportion of variants that are singleton is also the most strand biased for A>G/T>C variants, in the expected direction. “Coding” and “non-coding” indicate the strand of the labeled base change. In B and C, each base change pair (different base change on each strand) is labeled with the pair member where the ancestral base is A or C. Note: some pairs of points are completely overlapping at the resolution shown. D) Summary of overall average negative selection for 3’ UTR variants relative to intergenic variants (dashed line at 0), benchmarked against CDS variants that differentially affect encoded protein.

First, the vast majority of RBPs that bind predominantly in 3’ UTRs also bind coding sequence extensively (Van Nostrand et al. 2020), so calibrating to a transcribed region may dampen or obscure signals of negative selection from such regulatory elements. Relative to synonymous variants in transcribed coding regions, intergenic variants are under less selection (Karczewski et al. 2020) and are devoid of RNA regulatory elements characteristic of 3’ UTRs including RBP binding sites.

Second, there are many more intergenic variants available than synonymous coding variants (over 30-fold more, even after conservative filtering), providing greater statistical power to account for the influence of sequence (and other) contexts beyond dinucleotide composition that have been shown to substantially affect mutability (Carlson et al. 2018). Using intergenic variants, we were able to identify many such contexts that extended up to five bases on each side of the variant. This approach resulted in significantly different calibration across all base changes for the majority (54%) of all 3’ UTR variants, relative to a dinucleotide-only approach (Methods, Fig. 1B).

Third, existing work has seemingly performed calibration using paired dinucleotide contexts (e.g., reverse complements such as C[A>G]T and A[T>C]G are grouped together) (Lek et al. 2016; Karczewski et al. 2020; Whiffin et al. 2020). However, it is known that strand-biased processes including transcription-associated mutagenesis and transcription-coupled repair influence mutability in transcribed regions for some sequence contexts (e.g., A>G/T>C mutation rates are higher when A>G occurs on the coding strand; (Seplyarskiy et al. 2021; Seplyarskiy and Sunyaev 2021; Green et al. 2003), although this effect has not commonly been incorporated into mutation rate models (Carlson et al. 2018). To our knowledge, transcription-related bias has not yet been accounted for in any approach to calibrate negative selection analyses. We assessed the influence of transcription on mutability in 3’ UTRs using *de novo* mutation data (Turner et al. 2017) in addition to allele frequency data from gnomAD, where increased mutability presents as a decrease in the proportion of variants that are singletons. Consistent with previous work (Seplyarskiy et al. 2021), such effects were substantial for A>G/T>C base changes and minimal for other contexts (Fig. 1C). Variable rates of germline transcription across genes and across developmental contexts make it difficult to account for the influence of transcription in any calibration approach. Therefore, we excluded A>G/T>C variants from our analysis, where transcriptional effects are most apparent.

In addressing these issues, we have developed the intergenic MAPS or “iMAPS” approach, an extension of the MAPS metric (Lek et al. 2016; Karczewski et al. 2020), to quantify negative selection in non-coding regions including 3’ UTRs. We benchmarked 3’ UTR iMAPS values against canonical classes of coding variation. As a whole, negative selection in 3’ UTRs was higher than in intergenic regions and slightly lower than for synonymous coding variants, paralleling the pattern of evolutionary conservation across mammals in these regions (Lindblad-Toh et al. 2011). More specifically, we looked at regulatory element annotations in 3’ UTRs including eCLIP peaks marking RBP binding sites (Van Nostrand et al. 2020), miRNA target sites predicted by TargetScan+ (all targets of miRNA families conserved in mammals or more broadly) (Agarwal et al. 2015), and the canonical polyadenylation signal hexamer AWUAAA (W = A or U). In aggregate, none of these individual annotations had iMAPS scores exceeding that of synonymous variants (Fig. 1D), suggesting that this conventional approach of one-dimensional annotation intersection was insufficient to enrich for variants under negative selection and that variant interpretation in 3’ UTRs may require more nuanced approaches. We reasoned that there are subsets of variants within these annotations under strong selection, and hypothesized that further stratification of variants aided by additional annotations could uncover these classes of deleterious variants in 3’ UTRs.

We first focused on general RBP binding sites in 3’ UTRs, with the goal of identifying precise/short RBP binding sites of high confidence using complementary orthogonal methods: for each RBP with available data, we identified the highest affinity RBPamp motif (based on high throughput *in vitro* RNA Bind-n-Seq (RBNS) data (Dominguez et al. 2018; Jens et al. 2022) in the vicinity of each of the more than 25,000 ENCODE eCLIP peaks (Van Nostrand et al. 2020) and termed these sites “RBPamp eCLIP-Proximal” or “ReP” sites (Fig. 2A). At either 10 or 11 bases long, ReP sites were about five-times more precise than typical eCLIP peaks. The significant enrichment of ReP sites around the 5’ ends of eCLIP peaks highlights the coherence of these datasets and validates this approach to identify precise RBP binding sites marked by eCLIP peaks for the vast majority of available RBPs (Supplementary Fig. 1). Furthermore, we found that variants in ReP sites were up to 6-fold enriched for variants that altered transcript levels in a 3’ UTR massively parallel reporter assay (MPRA; (Griesemer et al. 2021) Fig. 2B). Together, these data suggest many ReP sites are RNA elements with endogenous RBP-binding and regulatory capacities.

**Fig. 2:**
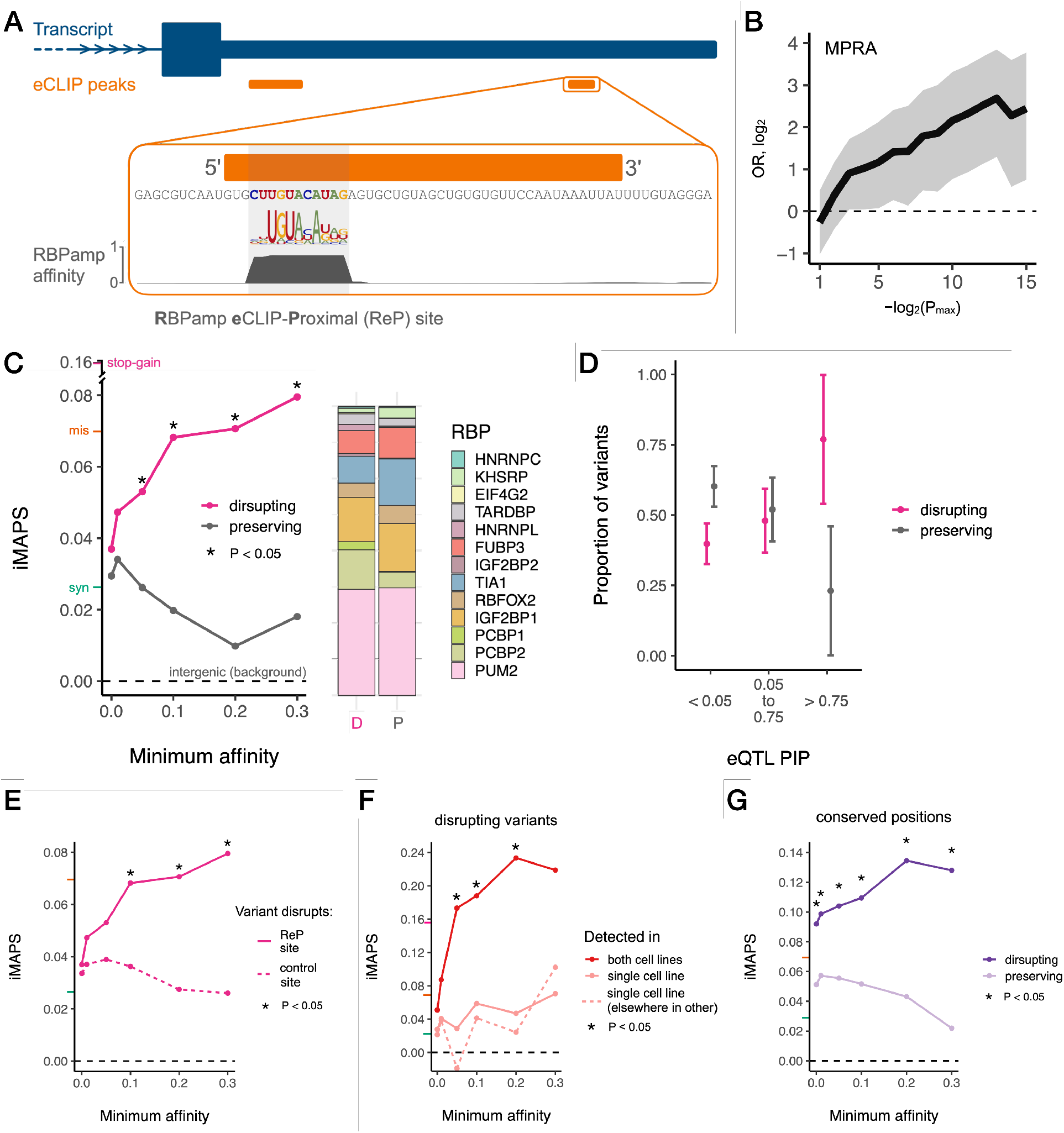
Variants disrupting ReP sites in 3’ UTRs are under strong negative selection. A) A ReP site is the highest affinity RBP binding site at an eCLIP peak for that RBP. B) ReP site variants are enriched for significant transcript abundance modulating activity in a massively parallel reporter assay of 3’ UTR variants (Griesemer et al. 2021). Shaded area indicates 95% confidence intervals. OR = odds ratio. P_max_ = P value significance threshold. C) Variants disrupting RBP affinity / binding at ReP sites are under stronger selection than those that preserve affinity / binding. The stacked bar plot shows the proportion of variants found within each RBP’s ReP sites. “D” = disrupting, “P” = preserving. iMAPS scores for synonymous (green), missense (orange), and/or stop-gain (pink) coding variants are shown on y-axis here and in E-G for reference. D) High PIP (likely causal) eQTL variants in ReP sites are much more likely to be disrupting than preserving. Error bars indicate 95% confidence intervals for different PIP ranges. E) Variants disrupting ReP sites are under stronger selection than those disrupting the control site with closest affinity in each gene. The dashed line at 0 indicates background levels of intergenic selection. F) Variants disrupting ReP sites detected by eCLIP in both cell lines are under stronger selection than variants disrupting ReP sites detected in only one cell line. Dashed line is data for a control set ReP sites detected in both cell lines but at different positions in the same 3’ UTR. G) At highly conserved positions (PhastCons 100-way = 1.0), variants disrupting ReP sites are under stronger selection than those that preserve RBP affinity/binding. The number of variants available for iMAPS analyses in C, and E-G are included in Supplementary Table 1.

Next, we leveraged RBPamp affinity models to predict the impact of each gnomAD variant within ReP sites on RBP binding. Variants where the derived allele was predicted to have substantially reduced affinity to the RBP relative to the ancestral allele were termed “disrupting,” while variants where the derived and ancestral alleles were predicted to have similar affinity were termed “preserving” (Methods). Here, we emphasized ReP sites that we term “focal,” where most of the local affinity to the RBP is concentrated in the ReP site itself. We reasoned that variants in focal sites would be more likely to have large and predictable impacts on regulation relative to variants in sites where the affinity is spread across tens of nucleotides, since the binding location can be more confidently inferred, and there is less potential for redundant regulation when affinity is concentrated to one short element. Indeed, for 3’ UTR ReP site variants included in an MPRA measuring transcript levels (Griesemer et al. 2021), variants in focal ReP sites conferred significantly larger alterations in transcript levels relative to variants in non-focal ReP sites (Supplementary Fig. 2).

Both disrupting and preserving variants were identified in ReP sites spanning over a dozen diverse RBPs with established regulatory roles in 3’ UTRs (Fig. 2C). We observed that disrupting variants were subject to significantly stronger negative selection than preserving variants, supporting the importance of this distinction. Furthermore, disrupting variants in increasingly high affinity ReP sites were under increasingly stronger negative selection. Above the highest minimum affinity analyzed (30% of an ideal binding site), the negative selection experienced by these variants exceeded the genome-wide average of missense coding variants (Fig. 2C). Conversely, for all minimum affinities analyzed, preserving variants did not surpass the much lower average negative selection experienced by synonymous coding variants (Fig. 2C), suggesting that such variants are similarly neutral to overall 3’UTR variants. Using orthogonal eQTL data, we also found that ReP site variants with increasingly higher PIPs (probability of causing an expression difference) were more likely to be disrupting than preserving (Fig. 2D). Collectively, these data demonstrate that RBPamp can be used to prioritize RBP binding site variants more likely to drive deleterious gene expression changes.

We considered whether biases related to gene expression might contribute to the above results. Since statistical power for eCLIP peak-calling in each gene is a function of eCLIP read counts for that gene, eCLIP peaks and therefore ReP sites are enriched in relatively highly expressed genes (Supplementary Fig. 3), which tend to be more constrained. For example, we found coding variants from genes with 3’ UTR ReP sites to have slightly elevated synonymous and missense iMAPS (~ 0.034 and 0.10, respectively) relative to genome-wide averages. However, when considering a set of control sites matched for gene expression and RBP affinity with ReP sites, we found that variants that disrupt ReP sites had substantially higher iMAPS scores than variants that disrupted these control sites (Fig. 2E). This analysis shows that the presence of eCLIP signal at ReP sites is an important feature in predicting constraint, and that independent of eCLIP, high RBP affinity and high gene expression are not generally sufficient to identify sites under strong selection.

In an effort to better understand other aspects of RBP binding sites in the context of negative selection, we focused on RBP binding events shared across cell lines, and on cross-species sequence conservation. Since eCLIP was performed in two distinct cell lines for many RBPs, we reasoned that ReP sites identified in both cell lines would be enriched for RBP binding sites that are broadly utilized across cell types and might therefore be involved in regulation of fundamental cellular processes. Indeed, we found that variants disrupting ReP sites present in both cell lines were under very strong negative selection (Fig. 2F). This was not simply a result of 3’ UTR eCLIP peaks detected in both cell lines deriving from more highly expressed genes (where increased read depth provides increased statistical power to call peaks), as disruption of a control set of ReP sites from genes where ReP sites were detected in both cell lines but at distinct 3’ UTR positions was much less deleterious. These results suggest that detection of eCLIP signal across distinct cell types enriches for binding sites that are functional.

When exclusively examining positions conserved across vertebrates, variants disrupting ReP sites were highly deleterious. Importantly, variants at the same sites that preserve RBP binding were under much weaker selection, supporting that preservation of RBP binding is a major contributor to the high conservation observed at these positions (Fig. 2G). This finding also highlights the capacity of our approach — unlike most measures of cross-species sequence conservation — to capture divergent selection signals based on the nature of the base change introduced by the variant. Overall, these results demonstrate that ReP sites, combining in-cell eCLIP and *in vitro* RBNS/RBPamp binding site data, reveal precise and functional regulatory sites bound by RBPs.

Moving beyond genetic variants within ancestral binding sites, we next explored whether derived variant alleles that create new RBP binding sites experience substantial selective pressure. We focused on creation or strengthening of Pumilio family (PUM1 and PUM2) binding sites since these RBPs bind with high affinity (sub-nM Kd) and can strongly reduce target gene expression via transcript destabilization (Etten et al. 2012). Importantly, since this and all subsequent analyses involve regulatory elements predicted based on primary sequence, we were attentive to the alternative polyadenylation (APA) context. Specifically, we distinguished between variants in “core” and “variable” 3’ UTR regions for each gene. Core regions are upstream of the most utilized or “primary” poly(A) site according to the average reads per million across all tissues and cell lines in polyA_DB (Wang et al. 2018). Variable regions are downstream of the primary poly(A) and are included in processed transcripts only when a less frequently selected poly(A) site is utilized (Fig. 3A). Applying this approach to the creation of RBP binding sites, we found that variants increasing Pumilio affinity are increasingly deleterious at higher derived allele affinities, and creation of sites at least 35% as strong as an ideal Pumilio site approached the strength of selection seen for missense coding variants. This strong signal was specific to the creation of binding sites within core 3’ UTR regions. Destabilization in core regions is expected to impact the most transcripts resulting from APA. Conversely, variants creating binding sites within variable 3’ UTR regions did not exceed levels of selection seen for synonymous coding variants (Fig. 3B). Using independent eQTL data, we verified that variants creating high affinity Pumilio binding sites had higher probabilities of being causal of gene expression differences (Fig. 3C) and were more likely to decrease gene expression (Fig. 3D). Thus, in addition to disruption of RBP binding being deleterious, creation of strong RBP binding sites can cause detectable levels of negative selection, at least in some contexts.

**Fig. 3:**
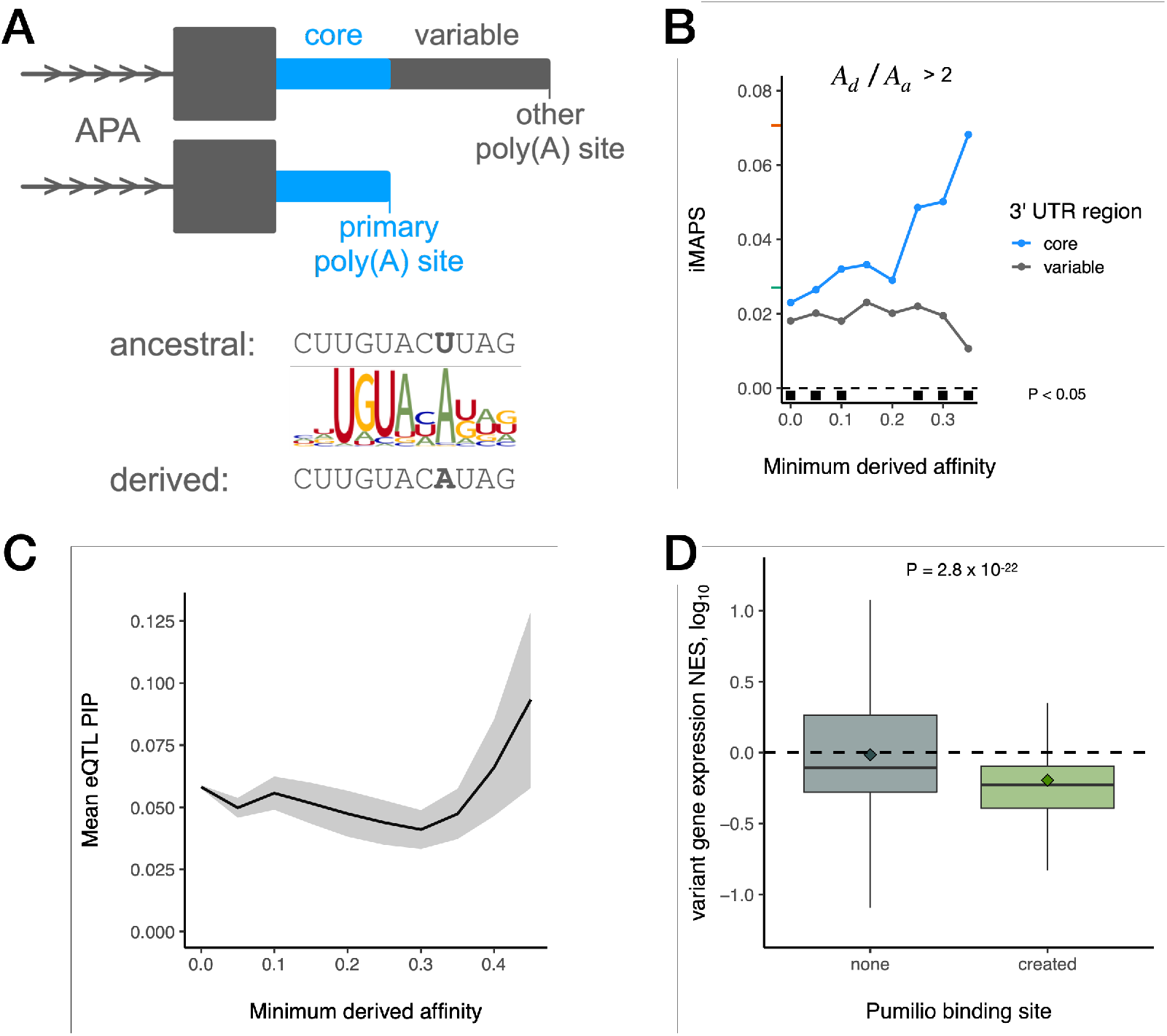
The impact of de novo Pumilio binding sites created by genetic variants. A) Top: Illustration of the “core” 3’ UTR region (blue) upstream of the primary (most utilized) poly(A) site and the “variable” 3’ UTR region (gray) present only in isoforms resulting from use of secondary poly(A) sites. Bottom: Schematic illustrating an example of Pumilio binding site creation. B) Variants creating Pumilio binding sites in core 3’ UTR regions are under strong negative selection. A_d_ = derived affinity, A_a_ = ancestral affinity. The number of variants analyzed are included in Supplementary Table 1. C) eQTL variants that create high-affinity Pumilio sites typically have higher PIP values. Shaded area indicates mean +/−standard error. D) eQTL variants creating Pumilio sites (n = 448 eQTLs) are also associated with decreased transcript levels relative to those with no site (n = 70,318 eQTLs), consistent with the destabilizing impact of Pumilio binding. NES = log_10_ normalized effect size. Diamonds indicate mean NES values for each category.

Since regulation by miRNAs is a widespread form of post-transcriptional gene regulation, we investigated negative selection at miRNA target sites (8mer, 7mer-m8, and 7mer-A1) in 3’ UTRs according to TargetScan (Agarwal et al. 2015). Targets of both “broadly conserved” (vertebrate-wide) and “conserved” (mammalian-wide) miRNA families were under stronger negative selection in core 3’ UTR regions upstream of primary polyadenylation sites compared to variable regions less often included in processed transcripts. Targets of more broadly conserved miRNAs were generally under stronger selection (Fig. 4A), consistent with the finding that there are fewer conserved targets of miRNAs conserved in mammals and not more broadly (Friedman et al. 2009). As a positive control, conserved target sites of these miRNA families were under stronger selection than non-conserved sites, as has been previously observed (Chen and Rajewsky 2006), exceeding average synonymous coding levels (Supplementary Fig. 4). We next sought to test whether certain subsets of miRNA target sites were under stronger selection, without relying on target conservation across species. We focused on the most potent target sites: 8-mer targets of broadly conserved miRNA families. Within these sites, we observed a positive association between AU-rich dinucleotides flanking target sites and negative selection (Fig. 4B). The regions flanking miRNA targets are enriched for Adenosine (Lewis et al. 2005), and targets in AU-rich contexts are known to confer stronger binding and increased repression, likely due to reduced secondary structure increasing target accessibility (Grimson et al. 2007; McGeary et al. 2019; Nielsen et al. 2007). We also found that overlapping targets of more than one miRNA family were under stronger negative selection (Fig. 4C), presumably because such overlap increases the odds that the site has regulatory activity.

**Fig. 4:**
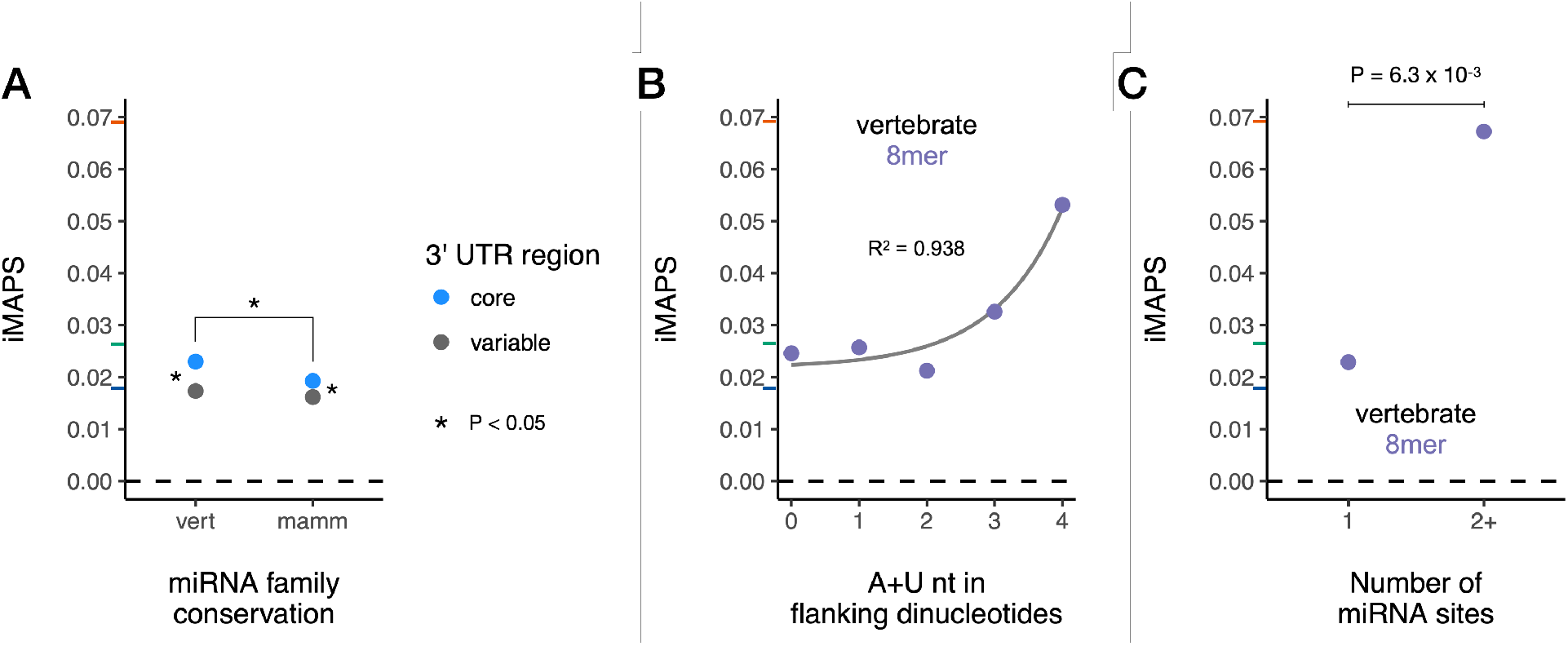
Negative selection at miRNA target sites in 3’ UTRs. A) miRNA targets (including 8mer, 7mer-m8, and 7mer-A1 target types) are under selection in core 3’ UTR regions for both broadly conserved (“vert” = vertebrate) and conserved (“mamm” = mammalian) microRNA families. n = 179,220 variants for vertebrate core, 197,573 for mammalian core, 137,103 for vertebrate variable, and 147,679 for mammalian variable. B) Stronger selection is observed for 8mer targets with increasing AU content of the dinucleotides immediately preceding and following the target site. The gray line is a linear model describing the data using an exponential fit. R^2^ shows Pearson’s product-moment correlation. C) Variants within targets recognized by two or more broadly conserved miRNA families are under stronger selection than those within targets of a single miRNA family. n = 10,912 variants for single targets and n = 389 variants for variants in targets of multiple miRNAs. y-axis tick marks indicate the genome-wide average iMAPS for 3’ UTR (blue), synonymous (green), and missense (orange) coding variants. The dashed line at 0 indicates background levels of intergenic selection.

To assess negative selection associated with polyadenylation, we focused on on the polyadenylation signal (PAS), as it is the major determinant of poly(A) site selection (Zhu et al. 2018; Hamilton et al. 2019; Shulman and Elkon 2020; Tian and Graber 2012). We focused specifically on variants in the two “top” PAS hexamers matching AWUAAA. We used data from the polyA_DB database (Wang et al. 2018) to classify AWUAAA hexamers as “primary” (30 to 15 bases upstream of the most utilized poly(A) site in a gene) or “secondary” (at any other 3’ UTR position). Analogous to our analysis of RBP motifs, we classified variants as “disrupting” if they resulted in loss of an AWUAAA motif without creating any other enriched hexamer from (Ni et al. 2013), with all other variants classified as “preserving” (Fig. 5A).

**Fig. 5:**
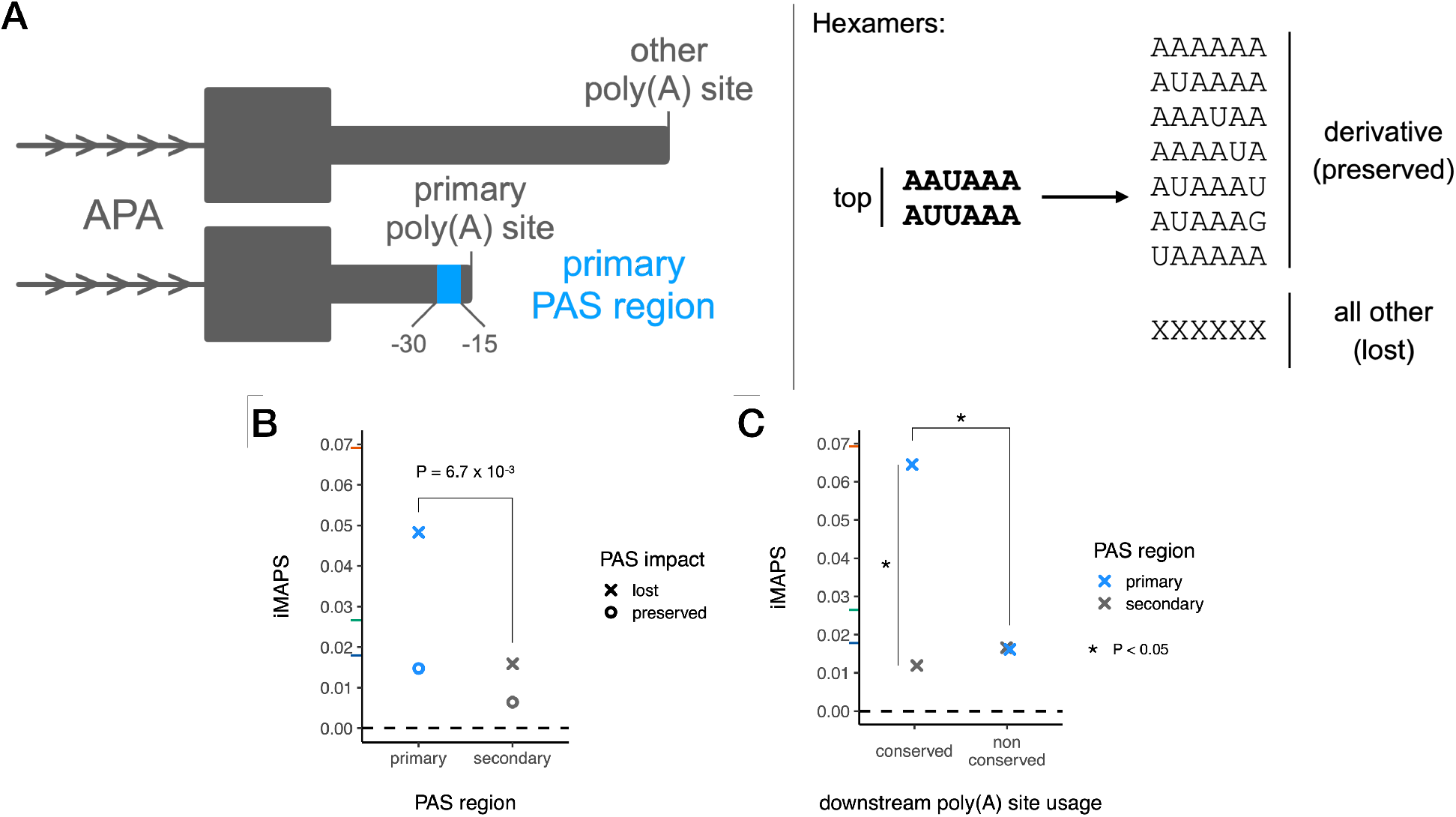
Negative selection of polyadenylation signals (PASs) in 3’ UTRs. A) Schematic of polyadenylation in 3’ UTRs. Left: the primary PAS region for a given gene (associated with the most commonly used poly(A) site) is shown in blue. AWUAAA hexamers found in any other regions (gray) are considered secondary. Right: Canonical / top PAS hexamers match AWUAAA. A PAS is considered preserved if the variant results in a derivative PAS hexamer, otherwise it is considered lost. B) PAS loss in primary PAS regions is under stronger selection than PAS lost in secondary regions. n = 1,571 variants for lost PAS in primary regions and n = 14,867 variants for lost PAS in secondary regions. C) PAS loss in primary regions is under stronger selection when the downstream poly(A) site is also utilized in mouse or rat (conserved). n = 1,041 variants for primary conserved, 2,928 for secondary conserved, and 530 for primary nonconserved. y-axis tick marks indicate the genome-wide average iMAPS for 3’ UTR (blue), synonymous (green), and missense (orange) coding variants. The dashed line at 0 indicates background levels of intergenic selection.

We found that disrupting variants in primary PASs were under stronger selection than disrupting variants in secondary PASs. Conversely, preserving variants experienced selection similar to background 3’ UTR levels (Fig. 5B). We next assessed selection in the context of human poly(A) sites that have a homologous active poly(A) site in either mouse or rat that we term “conserved” (Wang et al. 2018). We found that, in aggregate, virtually all of the selection against variants disrupting primary PASs was attributable to PASs associated with these conserved poly(A) sites. Only background 3’ UTR levels of selection were observed for disruption of primary PASs associated with non-conserved poly(A) sites (Fig. 5C). Our results are consistent with a recent analysis arguing that most secondary poly(A) sites are non-adaptive (Xu and Zhang 2018), with some exceptions, of course (Mayr 2020).

Having identified classes of genetic variation under strong selection acting in different modes of post-transcriptional gene regulation in 3’ UTRs, we sought to summarize the frequency of such variants. This analysis was done in a manner that controlled for biases in gene expression between datasets such that different iMAPS results could be compared. We labeled variants belonging to classes with iMAPS ≥ 0.06 (approaching the genome-wide average for missense coding variants) “highly disruptive,” and those belonging to classes with iMAPS ≥ 0.03 (greater than the genome-wide average for synonymous coding variants) “moderately disruptive.” We use the term “disruptive” based on the strong inference that these variants disrupt molecular interactions with regulatory function, analogous to how missense variants disrupt protein coding. These labels will be helpful in efforts to interpret genetic variation associated with disease, but alone are insufficient as evidence of pathogenicity (Richards et al. 2015). We were able to label over 5,000 gnomAD variants in 3’ UTRs (> 1 out of every 2,000) as highly disruptive and almost 20,000 gnomAD variants in 3’ UTRs (> 1 out of every 600) as moderately disruptive (Fig. 6A, Supplementary Table 2). While most of our classes did not exceed missense level selection, this is not to suggest that there is an upper limit on selection in 3’ UTRs. For example, we have demonstrated that variants at conserved positions disrupting ReP sites are under very strong selection (Fig. 2G), but chose not to include classes based on cross-species conservation in this summary. In all, one quarter of protein-coding genes had at least one 3’ UTR variant labeled as highly disruptive, and half of protein-coding genes had at least one 3’ UTR variant labeled as moderately disruptive (Fig. 6B).

**Fig. 6:**
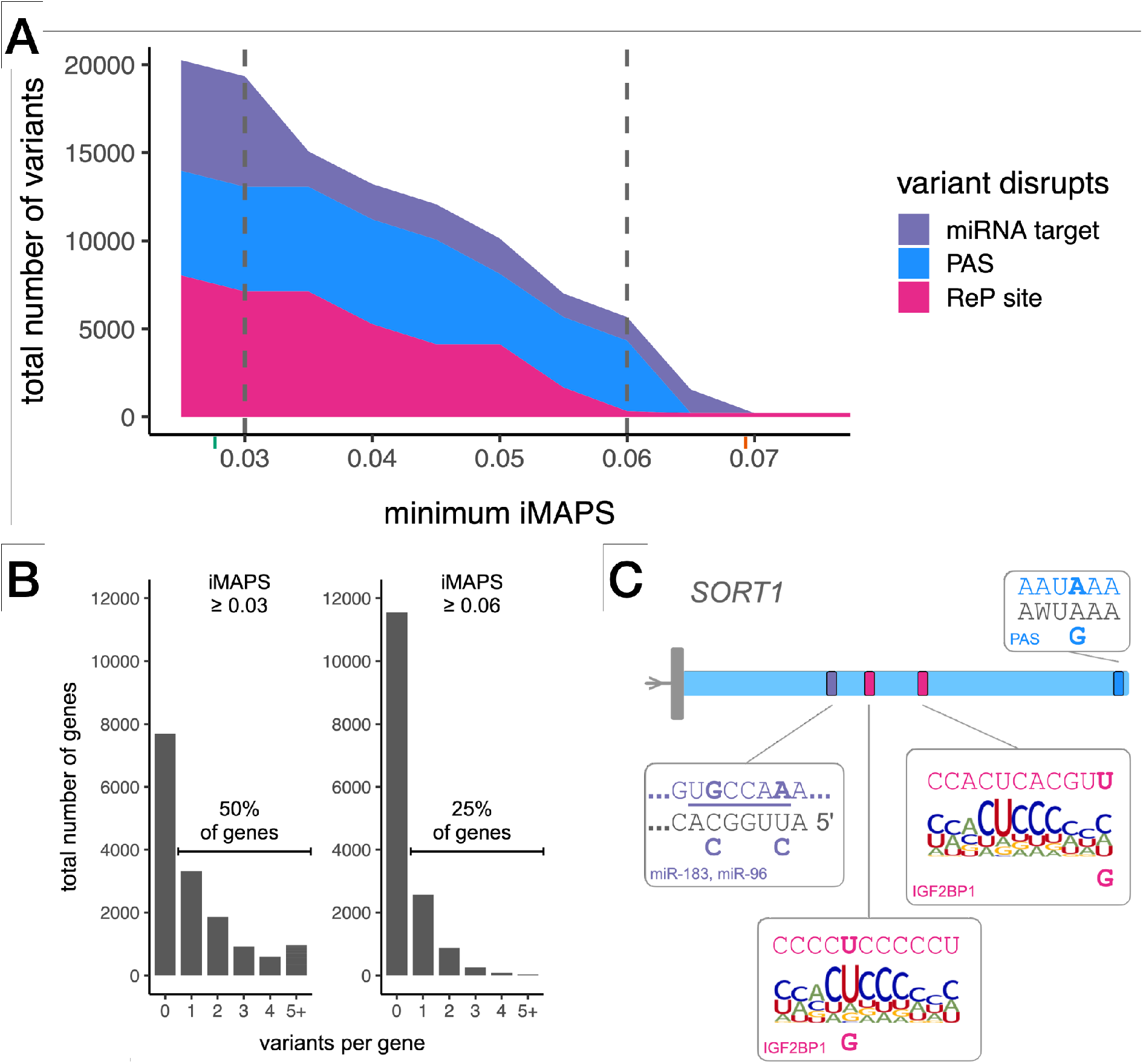
Summary of the number of gnomAD variants belonging to classes under strong selection in 3’ UTRs. A) The number of variants disrupting different regulatory elements in 3’ UTRs is shown for categories under increasingly strong selection (minimum iMAPS). x-axis tick marks indicate the genome-wide average iMAPS for synonymous (green) and missense (orange) coding variants. Vertical dashed lines indicate the cut-offs used for gene level analysis in B. B) Histogram of the number of disruptive variants per gene at two different thresholds. Left: half of all protein-coding genes contain at least one 3’ UTR variant labeled “moderately disruptive”. Right: one quarter of protein-coding genes contain at least one 3’ UTR variant labeled “highly disruptive”. C) Highly disruptive variants in *SORT1*. For clarity, only the 3’ UTR formed by the primary poly(A) site is shown. For each element, the reference sequence is shown at the top with ancestral variant alleles in bold, and alternative alleles are shown at the bottom. Mature miR sequence and top PAS hexamers are shown in gray and the miR seed region is underlined. ReP site variants are shown in the context of an RBPamp affinity model for IGF2BP1.

The data in Supplementary Table 2 can be used to augment conventional approaches for 3’ UTR variant interpretation for many human genes. As examples, we highlight two genes with known disease associations and highly disruptive 3’ UTR variants: insulin like growth factor 2 receptor (*IGF2R*) (Supplementary Fig. 5) and sortilin 1 (*SORT1*) (Fig 6C). *SORT1* encodes a sortilin family receptor involved in intracellular trafficking and is associated with cholesterol levels and myocardial infarction risk (Kjolby et al. 2015). *SORT1* is in the top 7% of loss-of-function-intolerant genes (rank 1,200), in the top 10% of genes by number of variants studied (896) (Chunn et al. 2020), and in the top 5% of genes by number of significant variant-trait GWAS associations (28) (Buniello et al. 2019). Five *SORT1* 3’ UTR gnomAD variants were labeled as highly disruptive: two disrupting an 8mer target of miR-183 and miR-96, two disrupting distinct IGF2BP1 ReP sites, and one disrupting the PAS for the primary (and conserved) poly(A) site (Fig. 6C). Notably, none of these types of annotations are reported by standard variant interpretation tools such as Variant Effect Predictor (McLaren 2016). In addition, while some (3/5) of these variants had high phyloP scores, a simple approach to identifying candidate regulatory variants by focusing on positions with phyloP ≥ 2 yielded over 17-fold more *SORT1* 3’ UTR gnomAD variants (a total of 87). As another example, *IGF2R* encodes a receptor for insulin-like growth factor 2 and is often mutated in hepatocellular carcinoma (De Souza et al. 1995). *IGF2R* is also a loss-of-function-intolerant gene where four 3’ UTR gnomAD variants were labeled as disruptive: three disrupting a PUM2 ReP site, and one disrupting the PAS for the primary (and conserved) poly(A) site (Supplementary Fig. 5). Again, there were over 26-fold more *IGF2R* 3’ UTR gnomAD variants at positions with phyloP ≥ 2 (a total of 107). By comparison, iMAPS-based labels are more specific, sensitive to variant base change and potential human-specific activity, and identify specific regulatory factors.

## Discussion

Recent efforts to catalog human genetic variation using deep sequencing on a genomewide basis and at the scale of tens of thousands of individuals have enabled study of the signatures of evolutionary selection in our genome (Karczewski et al. 2020). However, it has been difficult to detect signal within non-coding regulatory elements. Here we present the iMAPS method to more accurately and more confidently account for the impact of non-selective forces (such as mutability) on allele frequencies in non-coding regions of the transcriptome. Using iMAPS, we uncovered over 5,000 gnomAD variants belonging to classes under strong negative selection. These variants highly disrupt RBP binding, miRNA targeting, and cleavage/polyadenylation. Our approach classified variants using basic principles and data related to these modes of regulation, complementing efforts to interpret 3’ UTR variation using deep learning models (Park et al. 2021; Zhou et al. 2019). Layering internal controls for each set of analyses on top of our carefully controlled iMPAS metric lends confidence that the observed effects are driven by selection on variants affecting post-transcriptional gene regulation and not artifacts of forces independent of selection such as mutability. Collectively, to our knowledge, this work is the first to comprehensively and quantitatively assess negative selection at regulatory elements in human 3’ UTRs.

Many methods for variant interpretation (Rentzsch et al. 2018; Zhang et al. 2020) incorporate measures of conservation across species, as variants at highly conserved positions are generally more deleterious overall. However, reliance on cross-species conservation to identify functional genetic elements in non-coding regions has significant limitations. First, non-coding regions such as 3’ UTRs are of course less conserved than coding regions (Duret and Mouchiroud 2000; Lindblad-Toh et al. 2011). Furthermore, there is evidence that non-coding regulatory regions undergo more frequent lineage-specific adaptation, with the extent of lineage-specific constrained sequence rivaling or even exceeding that which is both constrained in humans and conserved across mammals (Ward and Kellis 2012; Rands et al. 2014). Here, we identify classes of 3’ UTR variants under negative selection in humans without relying on any measures of inter-species conservation, capturing human-specific regulatory constraints even at non-conserved positions. As an example of how constraint can differ within humans and between mammals, we found that variants disrupting PASs associated with less frequently used poly(A) sites in humans were not under strong selection, even in cases where the at poly(A) sites were also utilized in rat or mouse (Fig. 5C). A second limitation is that inter-species conservation scores such as PhastCons and PhyloP do not provide any information as to what extent possible alternative alleles may be differentially tolerated at a given position. We demonstrate the power of our analysis to differentially classify variants based on the consequence of the base change. This is seen most strikingly for RBP binding events where there are large differences in negative selection between disrupting and preserving variants within the same set of ReP sites, even when restricted to variants at the most highly conserved positions (Fig. 2G). Conventional one-dimensional approaches to variant interpretation typically prioritize all variants within an annotation. We show evidence that utilizing multi-dimensional annotations (e.g., ReP sites derived using both eCLIP and RBPamp data) is critical to uncovering classes of variation under strong selection, and that an approach sensitive to the impact of different potential base changes introduced by genetic variants can filter more neutral classes of variants within these annotations.

Our work uses orthogonal human population genetic data to validate the utility of several RNA related datasets for detecting regulatory elements in 3’ UTRs. We find the majority of selection against PAS-disruption, miRNA-disruption, and Pumilio binding site creation occurs in core 3’ UTR regions upstream of the most utilized poly(A) site in a gene, based on average utilization across various cell lines and tissues from polyA_DB (Wang et al. 2018), suggesting this consideration is extremely useful in guiding the search for functional elements in 3’ UTRs. Based on the notion that variants within expressed mRNA isoforms are more likely to be functional, we provide evidence that the APA context needs to be explicitly considered when interpreting 3’ UTR variants, as has proven useful in identifying targets of miRNAs (Nam et al. 2014). These results emphasize the utility of a recently developed “expression-aware” variant interpretation framework that considered how often variants were found within alternatively spliced transcript regions (Cummings et al. 2020), and argue for similar practices to be applied in 3’ UTRs based on APA. We also demonstrate the utility of performing eCLIP across multiple cell lines as RBP-disrupting variants within ReP sites supported by eCLIP peaks across two different cell lines are associated with the highest estimates of negative selection we observe in 3’ UTRs. Together, these results suggest that even though these datasets are derived from relatively arbitrary sets of mostly adult tissues and *ex vivo* cell lines, they are informative in uncovering selection that likely manifests in embryonic or other developmentally relevant tissues (Roux and Robinson-Rechavi 2008).

We also find that RBPamp affinity models (Jens et al. 2022) derived from *in vitro* RBNS data (Lambert et al. 2014; Dominguez et al. 2018) are insufficient to *independently* identify constrained RBP binding sites (Fig. 2E). However, when complemented by in-cell eCLIP data (Van Nostrand et al. 2020), these affinity models prove crucial for the identification of the precise binding sites associated with eCLIP peaks that are under selection. In describing these so-called ReP sites, we have introduced a powerful set of precise and high-confidence binding sites for a diverse set of RBPs. In addition to demonstrating that these sites are under negative selection, we also demonstrate they are enriched for regulatory function. We envision that ReP sites will serve as an annotation with utility for many applications and fill an important gap for variant interpretation in the context of general RBP binding. Moreover, at ReP sites, RBPamp is informative of a general minimum relative affinity that constitutes functional binding, and accurately distinguishes between variants that disrupt or preserve binding. Thus, when applied in the context of binding sites detected in cells, RBPamp affinity models derived fully from *in vitro* experiments are powerful tools to uncover functional binding events and to interpret variants that modulate RBP binding in human 3’ UTRs.

While using population genetic data from contemporary human populations to assess negative selection as presented here does not provide information at the level of individual genetic variants, our results can inform variant interpretation. The stratification of variants performed here provide useful thresholds and benchmarks for interpretation of both known and not yet observed variants that may impact different post-transcriptional gene regulatory mechanisms, enabling more confident interpretation of disease-associated non-coding variants (Fig. 6 and (Ellingford et al. 2022)). Specifically, we were able to label between 5,000 and 20,000 gnomAD variants in 3’ UTRs as moderately or highly disruptive, respectively, and highlight examples of haploinsufficient disease-associated genes with several highly disruptive 3’ UTR variants. While gnomAD is a useful reference, these numbers can be thought of as minima, as consideration of additional rare variants from other sources would increase the number of labeled variants. Overall, our findings deliver detailed insight into the nature of RBP activity in untranslated regions, helping us to continue to discover how these fundamental interactions regulate gene expression in ways that have and will continue to shape our biology as humans.

## Methods

### Genetic variants

Genetic variants from gnomAD release 3.0 were downloaded from https://gnomad.broadinstitute.org/downloads in vcf format. Single nucleotide variants passing gnomAD quality filters were isolated using VCFtools (Danecek et al. 2011). Genetic variants from denovo db (from non-SSC samples) were downloaded from https://denovo-db.gs.washington.edu/denovo-db/Download.jsp.

### Variant filtering for negative selection analyses

To reduce bias in the allele frequency spectra that might confound negative selection analyses, we filtered variants in: low complexity regions (defined by gnomAD), sex and mitochondrial chromosomes (X, Y, and M), regions with median sequencing coverage less than 25 (3/4 of overall median coverage) or greater than 42 (4/3 of overall median coverage), the ‘C’ position of CpG dinucleotides without any available methylation data, and CpG islands (Gardiner-Garden and Frommer 1987, Raney et al 2014). To ensure we were studying the impact of the newly derived allele, we excluded variants where the reference allele was not also identified as the ancestral allele using the *Homo sapiens* ancestor genome sequence from Ensembl (https://ftp.ensembl.org/pub/current_fasta/ancestral_alleles/homo_sapiens_ancestor_GRCh38.tar.gz). To limit the impact of transcription-associated mutability, all variants with base change of A>G on either strand (i.e. A>G or T>C) were excluded. Variants with any “N” base calls for the five nucleotides on either side of the variant were also excluded. Multi-allelic sites were retained and treated as independent variants. Variants in 3’ UTRs that were also within any Gencode CDS exons (on either strand) were excluded from analysis.

### Variant annotations

Variant positions were then annotated as either overlapping or non-overlapping with DNase hypersensitivity peaks (hg38wgEncodeRegDnaseClustered from the UCSC Table Browser), CpG islands (hg38cpgIslandExtUnmasked from the UCSC Table Browser), and H3K9me3 peaks (E062-H3K9me3.bed downloaded using https://github.com/carjed/smaug-genetics/blob/master/download_ref_data.sh). CpG transition variants were intersected with methylation data (processed data from Roadmap Epigenomics Consortium 2015; provided by gnomAD). These variants were assigned a methylation level as in (Karczewski et al. 2020): High = mean methylation > 60, medium = mean methylation ≤ 60 and > 20, low = mean methylation ≤ 20. Variants at positions with missing methylation data were not assigned a bin and were later excluded.

### Intergenic regions

The intergenic regions used to derive the expected proportion singleton values consisted of all regions at least 25 kb from any Gencode v32 entry. Gap locations and centromere regions were then excluded. To avoid disproportionate contributions from very large intergenic stretches in the genome, long continuous regions were trimmed to a maximum length of 1.5-times the 75^th^ percentile length. This resulted in a set of intergenic regions totaling approximately 330 Mb.

### Intergenic mutability-adjusted proportion singleton (iMAPS)

We used variants in the intergenic regions described above to create a calibration set of expected proportion singleton (variants only observed once in the sample) values for different genomic contexts. We considered nucleotide composition flanking the variant, CpG methylation levels, as well as overlap with DNase hypersensitivity peaks, H3K9me3 peaks, and CpG islands. Collectively, these annotations explain much of the variability observed in DNA mutability across different loci (Carlson et al. 2018). Since CpG islands had disproportionately large effect on mutability, and are very rare in 3’ UTRs, variants in CpG islands were simply excluded.

Next, a set of 416 contexts was generated as follows: 4^3^ = 64 dinucleotide contexts (including the 4 NCG>NTG contexts) × 3 base changes = 192 reverse complement pairs = 96 distinct contexts + 8 for NCG>NTG at two additional methylation levels = 104 × 2 (DNaseHS peak binary) = 208 × 2 (H3K9me3 peak binary) = 416. An example context is C[A>G]G, DNaseHS-positive, H3K9me3-negative. We then calculated a proportion singleton for each context as the number of singletons / total number of variants. Next, we considered an additional base of sequence context in each direction beyond the dinucleotide context in a stepwise fashion, generating all possible daughter contexts at the trinucleotide, tetranucleotide, pentanucleotide, and hexanucleotide levels for each parent dinucleotide context. At each level, we calculated the proportion singleton for intergenic variants. We then compared these daughter context proportion singleton values to the proportion singleton for the parent dinucleotide-level context using two-sided Fisher exact tests and Benjamini-Hochberg correction for multiple hypothesis testing. For each hexanucleotide-level context, we used the longest parent context (e.g. pentanucleotide over trinucleotide) with a proportion singleton significantly different than the dinucleotide-level context proportion singleton at an FDR of 0.2. For cases where there was no significant impact of considering additional sequence context, the dinucleotide-level context was used. These intergenic proportion singleton values were then assigned to each 3’ UTR variant with matching context, as the “expected singleton” value. For a given set 3’ UTR variants, intergenic mutability-adjusted proportion singleton (iMAPS) was calculated as the sum of observed singletons minus the sum of expected singletons, divided by the total number of variants.

To test for significant differences in negative selection between a group of variants we hypothesized to be under strong selection and a control group of variants, we performed one-sided Fisher exact tests comparing the number of observed singletons and non-singletons. Importantly, we tested against an odds ratio derived from comparing the number of expected singletons and non-singletons derived from the matched intergenic variants. Exact P values are reported for panels with a single statistical test. For those with more than one statistical test (for example testing at different minimum affinity thresholds), we used asterisks (*) to indicate tests with P < 0.05.

### Coding sequence (CDS)

Gencode v32 CDS exons were downloaded from the UCSC Table Browser. Only CDS exons with transcript ids matching Gencode v32 protein-coding transcripts from protein-coding genes were analyzed. Gencode phase information was used to determine codon positioning and protein-coding consequence. CDS exons with any amount of overlap with another CDS exon annotation with a different reading frame were excluded from analysis. Variants within three bases of any splice site were excluded from analysis. A very small number of variants where the ancestral allele coded for a stop codon were excluded from analysis.

### Human 3’ UTRs

Human 3’ UTRs were defined using polyA_DB version 3.2 (Wang et al. 2018). Each poly(A) site was matched to the closest upstream stop codon annotated in Gencode v32 to form a 3’ UTR. Only poly(A) sites downstream of annotated stop codons from protein-coding genes were considered for analysis. Resulting 3’ UTRs longer than 50 kb were filtered from analysis. For identification of ReP sites within 3’ UTRs, Gencode v32 3’ UTRs were also used for genes with no poly(A) site data in polyA_DB. For all analyses where core and variable 3’ UTR regions were stratified (see below), only polyA_DB inferred 3’ UTRs were used.

### RBPamp

Position-specific affinity matrices (PSAMs) of length k = 10 or k = 11 generated by RBPamp (Jens et al. 2022) were used to score the affinity of a given RBP for potential target RNA sequences relative to the RBP’s ideal binding site (affinity = 1.0). We used PSAMs that were generated by considering both the sequence and predicted secondary structure of random sequence target RNA oligos in RBNS experiments.

### RBPamp eCLIP-proximal (ReP) sites

eCLIP peak data for K562 and HepG2 cell lines were downloaded from the ENCODE data portal. Peaks that were reproducible (IDR) across biological replicates were used for analysis. RBPamp PSAMs were used to score the relative affinity of every possible binding site within 75 bases of the 5’ end of each eCLIP peak (in either direction). Reference sequence was used as input. We identified the highest affinity site associated with each eCLIP peak. We termed these sites “RBPamp eCLIP-proximal” or “ReP” sites and considered the union of ReP sites across both cell lines. Since eCLIP peaks are typically > 50 bases (and often hundreds of bases) in length, and the RBPamp models consider a maximum binding footprint of k = 10 or k = 11 bases (depending on the RBP), ReP sites are a more precise set of high-confidence RBP binding sites. For most RBP-cell line pairs, the highest scoring sites were highly enriched at or close to the 5’ end of eCLIP peaks. This is expected from eCLIP experiments, given that cross-linked RNA bases often interfere with reverse transcription, causing read pileup and subsequent peak calling at or downstream (3’) of cross-linking sites. Identification of the highest affinity sites was also used to validate RBPs for which eCLIP and RBPamp data were coherent: For the union of all peaks across cell lines for each RBP, we determine how frequently each position (+/−75 bases relative to the 5’ end of each peak) overlapped the highest affinity site. We then conducted a simulation where we select a random site for each peak (+/−75 bases from the 5’ end of the peak) with equal probability across all positions. We repeat this 10,000 times to obtain a P value of how often the simulated data results in a single position that is observed as frequently as or more frequently than the most frequently observed position for the real set of highest affinity sites. ReP sites for RBPs with P values < 0.01 were retained for negative selection analyses. This resulted in filtering of only a few RBPs, most with very few eCLIP peaks in 3’ UTRs (Supplementary Fig. 1).

Control sites used in Fig. 2E were generated by first masking all ReP sites so any site overlapping any part of a ReP site would receive an affinity score of 0. For each ReP site, we then identified the site in the same 3’ UTR with the closest affinity (either higher or lower). Variants in these sites were classified using the same method applied to variants in ReP sites as described below.

For RBPs with eCLIP data available for both K562 and HepG2 cells, ReP sites were considered shared across cell lines in Fig. 2F if the exact same ReP site was identified in each cell line for the same RBP, regardless of whether the associated eCLIP peaks matched exactly. Partially overlapping but not completely matching ReP sites were not considered shared across cell lines. Since ReP sites shared across cell lines required an eCLIP peak to be called twice (once in each cell line), these ReP sites may be enriched for highly expressed genes (that are more constrained in general) relative to ReP sites identified in only a single cell line. Therefore we used a control set of ReP sites from genes where 3’ UTR ReP sites were detected for the same RBP in both cell lines, but at different positions (i.e. they were not the exact same ReP site).

### MPRA transcript abundance modulating variants (tamVars)

TamVar data was obtained from (Griesemer et al. 2021). Variants with statistically significant tamVar activity in any cell line tested were considered significant. We tested the odds ratio (odds of ReP site variants having significant tamVar activity in the MPRA relative to all other variants tested) at decreasing P value cut-offs (increasing stringency).

### Classification of ReP site variants

ReP site variants were classified as either focal or non-focal and disrupting or preserving. First, for each variant allele, we summed the relative RBPamp affinities across all positions overlapping the variant (i.e. 11 affinities for a width k = 11 RBPamp model) and termed these the ancestral affinity and the derived affinity. Next, we summed the relative affinity values spanning from 25 bases upstream to 25 bases downstream of the ReP site and termed this the local affinity. Focal variants were those with ancestral affinity / local affinity > 2/3 (i.e. the majority of the local affinity could be attributed to the ReP site). Non-focal variants were those with ancestral affinity / local affinity < 1/3, or with another ReP site for the same RBP within 25 bases. Disrupting variants were those with derived affinity / ancestral affinity < 1/3, and preserving variants were those with derived affinity / ancestral affinity > 2/3. Negative selection analysis of disrupting and preserving variants was performed on variants aggregated across all RBPs with available data (after filtering as described above). Analysis of individual RBPs was not possible due to low statistical power.

### eQTLs

DAG-P fine-mapped eQTL variant call files were downloaded from the GTex Portal. For each variant, the maximum posterior inclusion probability (PIP) value across tissues was used. Due to the relatively small number of ReP site variants with fine-mapped eQTL data available, we used more relaxed cut-offs for defining disrupting (derived affinity / ancestral affinity < 1/2) and preserving variants (derived affinity / ancestral affinity > 1/2) for the analysis in Fig 2D. Notably, it is not possible to analyze negative selection acting on variants from association-based analyses such as GWAS and eQTL, as they are highly skewed toward common variants (with more statistical power to detect significant effects) and measures such as iMAPS depend on unbiased allele frequencies.

### Expression data

Gene expression summary data for HepG2 and K562 cell lines were downloaded from the ENCODE Portal. Genes with ids matching Gencode v32 protein-coding genes were analyzed. Mean transcripts per million values across replicates were used.

### Defining core and variable 3’ UTR regions

We used polyA_DB version 3.2 data to identify core and variable 3’ UTR regions (Wang et al. 2018). These regions were defined based on relative *utilization* of different poly(A) sites, not necessarily their relative position. We defined the primary poly(A) site for each gene as the site with the highest mean reads per million across all polyA_DB tissues and cell lines. Thus, our primary poly(A) sites capture the effects of both alternative last exon selection and alternative polyadenylation. All positions between the primary poly(A) site and its upstream stop codon, regardless of any intervening poly(A) sites, were considered core. All positions between the primary poly(A) site and the most distal poly(A) site were considered variable. If the most distal poly(A) site associated with a given stop codon was also the primary poly(A) site, we considered that 3’ UTR to have no variable region.

### Pumilio site creation

We used the RBPamp model for PUM1/PUM2 to predict the relative ancestral and derived affinities for all 3’ UTR gnomAD variants as described above. RBPamp was used to score relative affinity for PUM1/PUM2. Variants with derived affinity / ancestral affinity > 2 were included for negative selection analysis in Fig. 3B. All variants with alternative affinity > reference affinity were included for analysis of mean eQTL PIP. For eQTL gene expression analysis, the normalized effect size (NES) values obtained from GTEx represent the relative expression difference between transcripts containing the alternate allele and transcripts containing the reference allele, where positive values indicate higher expression for the alternate allele, and vice versa. eQTL variants with alternative affinity > 0.3 and alternative affinity > reference affinity were labeled as “created.” Variants labeled as “none” consisted of all other variants after excluding variants with reference affinity > 0.3 and those in PUM2 ReP sites. A Wilcoxon Rank Sum test was performed to compare NES values between the “none” and “created” groups. Each PIP-tissue combination was considered for each eQTL variant.

### miRNA target sites

All miRNA target site data was downloaded from https://www.targetscan.org/vert_80/vert_80_data_download/All_Target_Locations.hg19.bed.zip. Only 8mer, 7mer-m8, and 7mer-A1 target sites (including both conserved and non-conserved sites) of miRNA families conserved in mammals or vertebrates were considered, and targets of poorly conserved miRNA families were excluded. All considered targets were grouped together for analysis of miRNA targets in Fig. 1D. For all analyses variants in the following target positions were analyzed: bases within the seed pairing with positions 2 through 7 of the miRNA (all types), the base across from position 1 (for 8mer and 7mer-A1 types), and the base pairing with position 8 (8mer and 7mer-m8 types).

### Polyadenylation signal (PAS) analysis

“Top” PAS hexamers (with ancestral sequence AWUAAA, where W = A or U) 15 to 30 bases (inclusive) upstream of primary poly(A) sites identified from polyA_DB data (see above) were considered primary PASs. All other PASs (including those nearby any non-primary poly(A) sites in core regions) were considered secondary. PASs were considered lost if the derived allele sequence did not overlap any PAS hexamer, including both AWUAAA and other similar A-rich hexamers found to be enriched in PASs from (Ni et al. 2013). Variants with derived allele sequence that overlapped one or more PAS hexamers were considered preserved. Conserved and non-conserved (active in human and either rat or mouse) poly(A) site classifications were obtained from polyA_DB (Wang et al. 2018). To account for poly(A) sites that were difficult to confidently assign to a single stop codon/3’ UTR in the absence of long read sequencing data, we filtered pairs of 3’ UTRs where any poly(A) sites are within 100 bases of a downstream stop codon/3’ UTR start.

### Comparing iMAPS across modes of regulation

We scaled ReP site iMAPS values to facilitate comparison between all three modes of regulation investigated. ReP sites are derived from eCLIP peaks, which are enriched in highly expressed genes where higher read counts increase the statistical power for peak calling. Since highly expressed genes are more constrained, we expect genes containing ReP sites to have slightly elevated iMAPS relative to genome-wide averages. In contrast, miRNA targets were predicted and analyzed for all genes, and PASs were considered equally across all transcripts with a detectable 3’ end / poly(A) site.

We calculated iMAPS for synonymous and missense variants from genes with 3’ UTR ReP sites to scale ReP site variant iMAPS values. Specifically, we sampled CDS variants from genes with 3’ UTR ReP sites in a weighted fashion to reflect the number of ReP sites in each gene. These CDS variants were then classified and iMAPS was calculated as described above, resulting in iMAPS of ~ 0.034 and 0.10 for synonymous and missense variants, respectively. These values were used as relative benchmarks to convert ReP site iMAPS values to the genome-wide scale with synonymous and missense iMAPS of 0.028 and 0.070, respectively. For example, a ReP site iMAPS value of 0.067 (half-way between synonymous and missense ReP site gene values) would scale to 0.049 (half-way between synonymous and missense genome-wide values).

After scaling of ReP site iMAPS values as described above, we counted the number of gnomAD variants belonging to any 3’ UTR element-disrupting classification with iMAPS values above a minimum threshold ranging from 0.025 to 0.075 (in increments of 0.005). We chose 0.03 and 0.06 as minimum iMAPS thresholds to highlight as they exceeded genome-wide synonymous levels and approached genomewide missense variant levels, respectively. None of the filters described in “Variant filtering for negative selection analyses” above were applied when counting the number of gnomAD variants belonging to each classification.

## Supporting information

Supplementary Table 1: number of variants

Supplementary Table 2: disruptive 3' UTR variants

## Acknowledgments

We thank members of the Burge laboratory for their helpful discussions and comments on the manuscript, especially Marvin Jens and Hannah Jacobs for their insights. We thank Nicola Whiffin for helpful discussions and Konrad Karczewski for facilitating access to useful datasets. We also thank David Bartel and Evan Boyle for providing helpful comments on the manuscript. S.D.F. was supported by a postdoctoral fellowship from the Natural Sciences and Engineering Research Council of Canada (NSERC). This work was funded by grants from the NIH (GM085319 and HG002439 to C.B.B.).

## Author Contributions

S.D.F. designed the study with input from C.B.B; S.D.F. performed analysis with contributions from L.R.; S.D.F. wrote the draft manuscript; S.D.F. and C.B.B. finalized the manuscript with input from L.R.

