## Supplementary figures 1-5 for "Quantifying negative selection in human 3’ UTRs uncovers constrained targets of RNA-binding proteins"

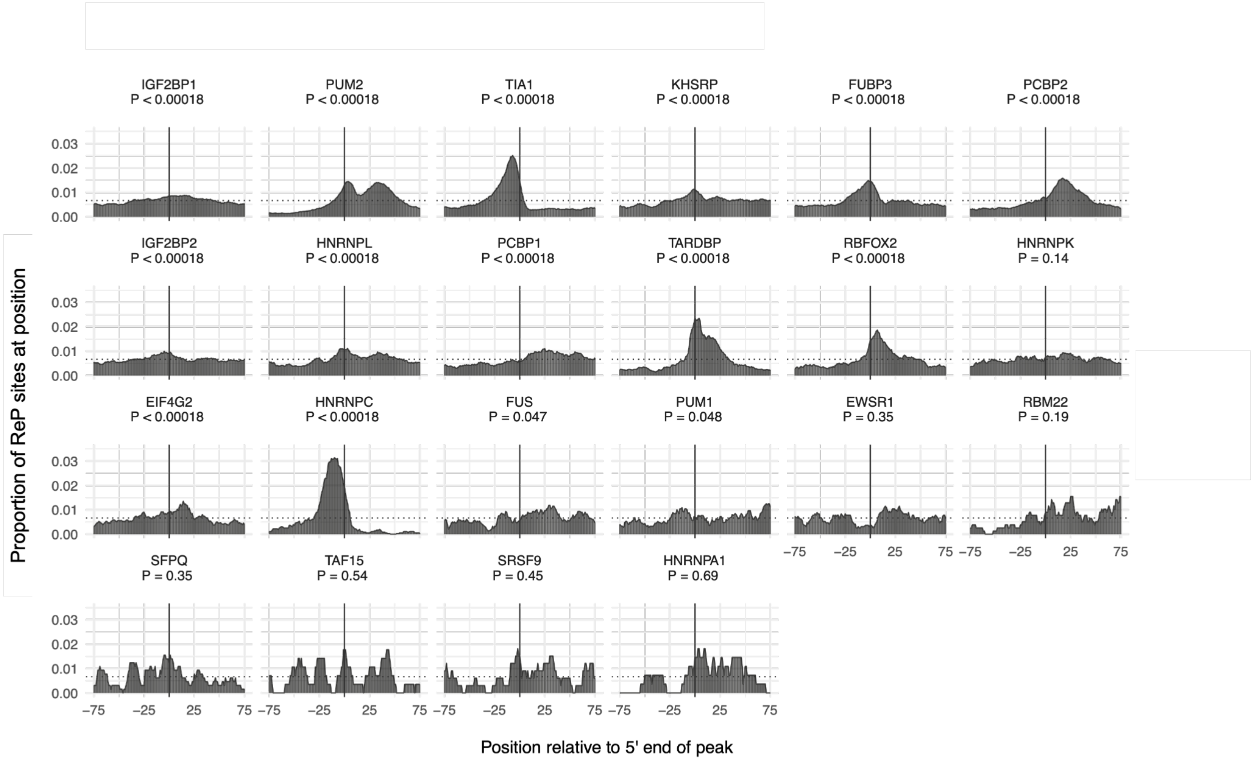


**Supplementary Fig. 1:** ReP sites are typically located close to the 5' end of eCLIP peaks in 3' UTRs. The union of eCLIP peaks across both HepG2 and K562 cell lines was used for RBPs with clip data in both cell lines. RBPs are sorted in order of descending number of peaks in 3' UTRs. Vertical lines indicate eCLIP peak 5' ends. The dashed horizontal lines indicate the null expectation of uniformly distributed highest affinity sites relative to eCLIP peaks. P values represent the proportion of 10,000 simulations with a maximum proportion value greater than the actual observed value, after Benjamini-Hochberg adjustment for multiple hypothesis testing.

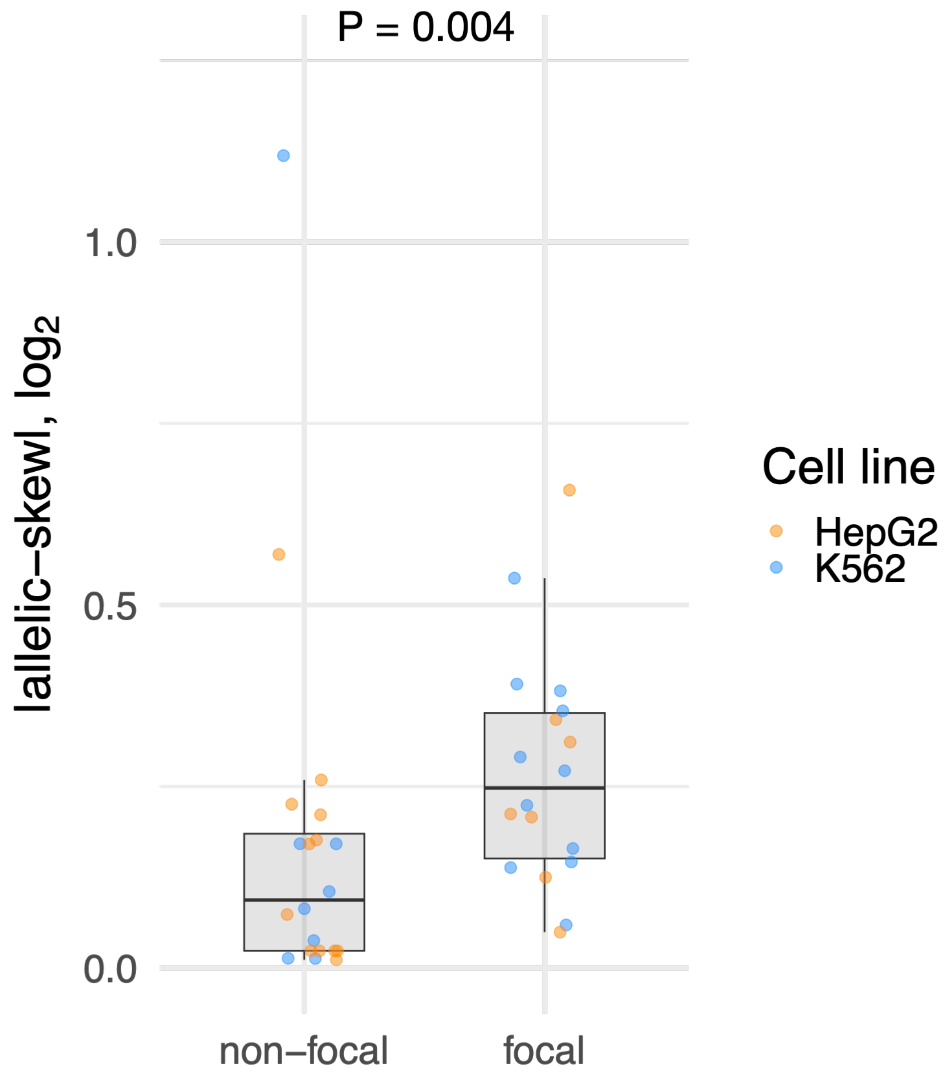


**Supplementary Fig. 2:** Focal ReP site variants promoted significantly larger skews in steady-state transcript levels than non-focal ReP site variants in a 3' UTR MPRA. P value is the result of a Wilcoxon rank sum test with continuity correction.

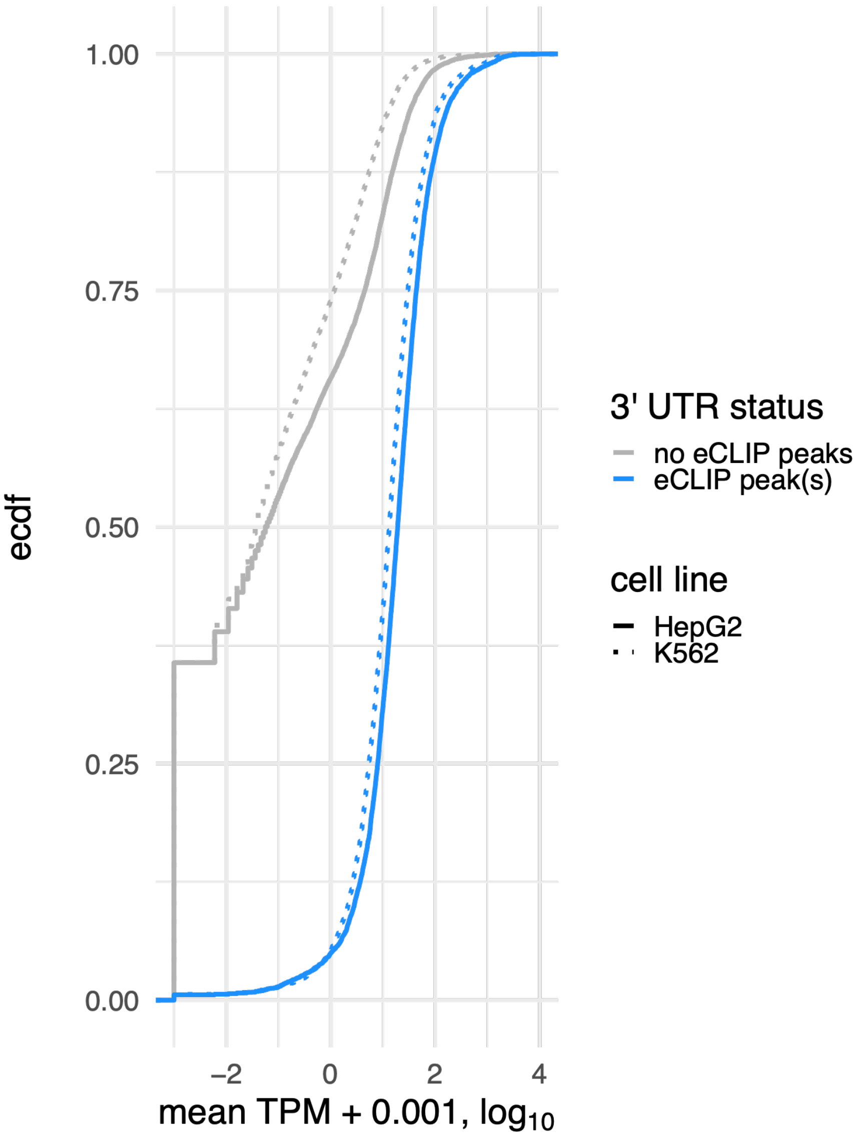


**Supplementary Fig. 3:** Transcripts with 3' UTR eCLIP peaks (for any RBP) are more highly expressed in HepG2 and K562 cells. TPM = transcripts per million. Values shown are means across two replicates for each cell line.

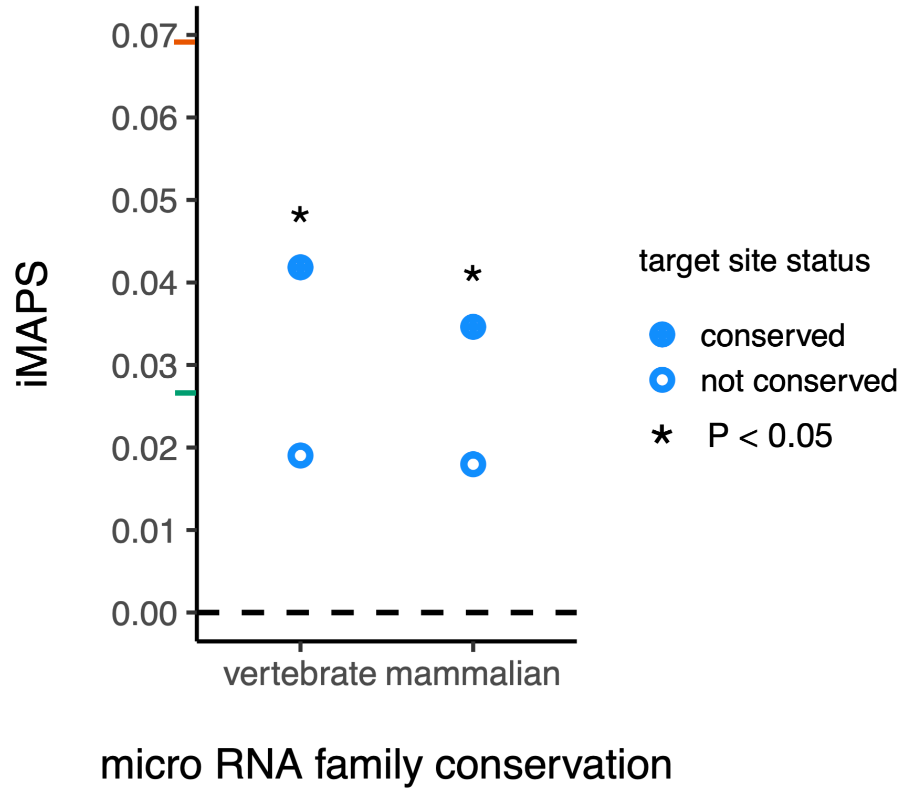


**Supplementary Fig. 4:** Conserved miRNA targets are under stronger selection than non-conserved targets. P values are the result of one-sided Fisher-exact tests. iMAPS scores for synonymous (green) and missense (orange) coding variants are shown on y-axis for reference.

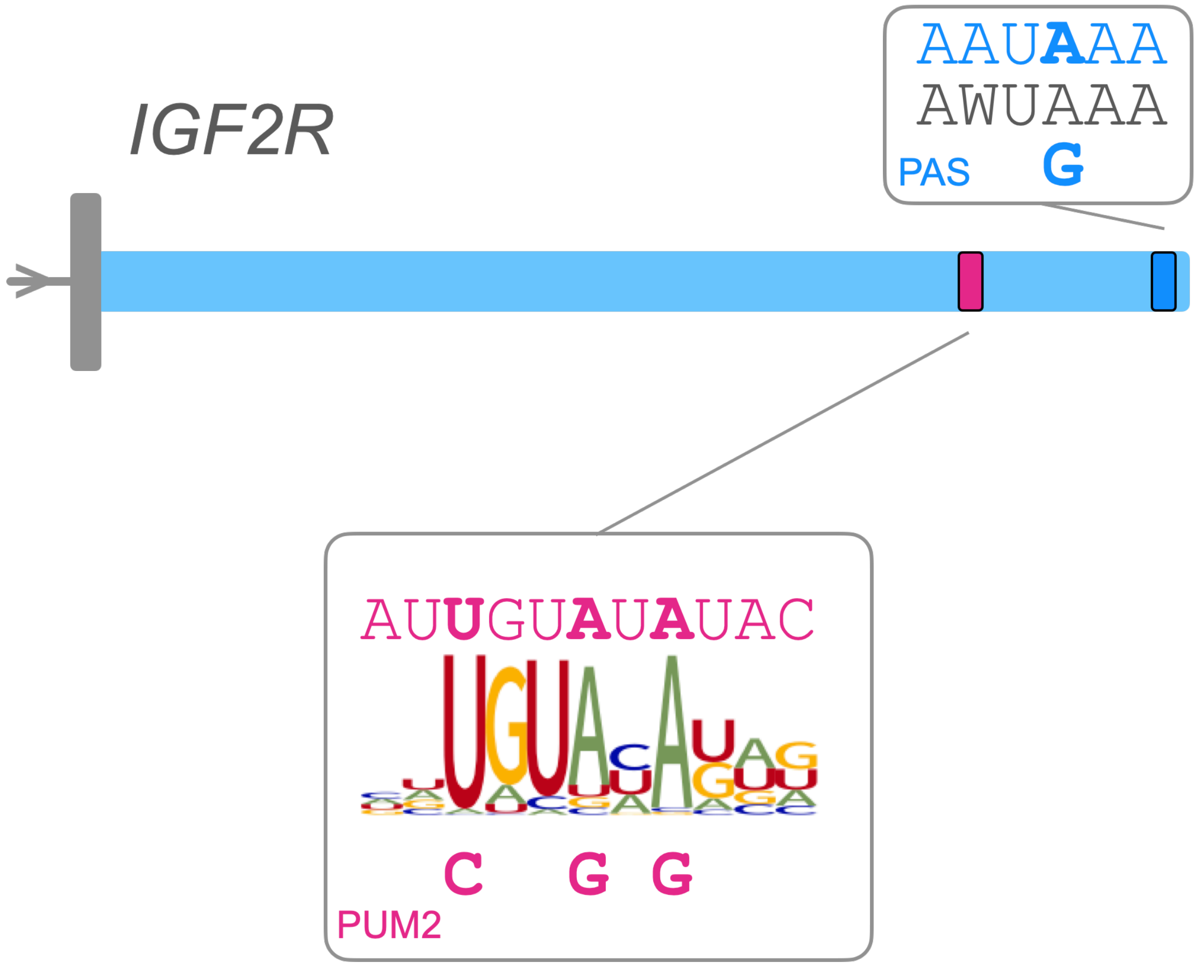


**Supplementary Fig. 5:** Highly disruptive variants in the *IGF2R* 3' UTR. Four 3' UTR gnomAD variants were labeled as disruptive*: three disrupting a single PUM2 ReP site, and one disrupting the PAS for the primary (and conserved) poly(A) site. For each element, the reference sequence is shown on the top with ancestral alleles in bold. Derived alleles are shown on the bottom. ReP sites are shown in pink, and PAS in blue. The top PAS hexamers are shown in gray. ReP site variants are shown in the context of an RBPamp affinity model for PUM2. *Note that the ReP site variants belong to a class with a minimum iMAPS of 0.05, while the PAS variant belongs to a class with a minimum iMAPS of 0.06.
